# FEATURE-SPECIFIC ANTICIPATORY PROCESSING FADES DURING HUMAN SLEEP

**DOI:** 10.1101/2024.09.16.613294

**Authors:** Pavlos I. Topalidis, Lisa Reisinger, Juliane Shubert, Mohamed S. Ameen, Nathan Weisz, Manuel Schabus

**Affiliations:** Centre for Cognitive Neuroscience and Department of Psychology, University of Salzburg, Austria; Neuroscience Institute, Christian Doppler University Hospital, Paracelsus Medical University, Salzburg, Austria

## Abstract

Imagine you are listening to a familiar song on the radio. As the melody and rhythm unfold, you can often anticipate the next note or beat, even before it plays. This ability demonstrates the brain’s capacity to extract statistical regularities from sensory input and to generate predictions about future sensory events. It is considered automatic, requiring no conscious effort or attentional resources (1–4). But to what extent does this predictive ability operate when our attention is greatly reduced, such as during sleep? Experimental findings from animal and human studies reveal a complex picture of how the brain engages in predictive processing during sleep (5–13). Although evidence suggests that the brain differentially reacts to unexpected stimuli and rhythmic music (5,7,13), there is a notable disruption in feedback processing, which is essential for generating accurate predictions of upcoming stimuli (10). Here, for the first time, we examine the brain’s ability during sleep to predict or pre-activate low-level features of expected stimuli before presentation. We use sequences of predictable or unpredictable/random tones in a passive-listening paradigm while recording simultaneous electroencephalography (EEG) and magnetoencephalography (MEG) during wakefulness and sleep. We found that during wakefulness, N1 sleep and N2 sleep, subtle changes in tone frequencies elicit unique/distinct neural activations. However, these activations are less distinct and less sustained during sleep than during wakefulness. Critically, replicating previous work in wakefulness (4), we find evidence that neural activations specific to the anticipated tone occur before its presentation. Extending previous findings, we show that such predictive neural patterns fade as individuals fall into sleep.

**In Brief:** The extent to which predictive processing takes place in sleep is yet to be determined. Using a passive-listening EEG/MEG paradigm, Topalidis et al. show that auditory representations in sleep are brief and unstable, easily overwritten by subsequent inputs, which possibly hinders the tracking and extraction of sensory associations.

**Highlights:**

- Participants passively listened to random and predictable sequences of tones during both wakefulness and sleep, without being made aware of the underlying pattern.
- The brain reta
- ins the ability to process basic low-level features during sleep.
- While these feature-specific responses are preserved during sleep, they are less distinct and sustained than in wakefulness.
- Unlike in wakefulness, during sleep, the brain does not predict or anticipate upcoming sounds, despite continuing to process basic auditory information.

## RESULTS

Since the study of predictive processes during sleep has been limited to looking at how the brain reacts to an unexpected stimulus (compared to an expected one) (14), we aimed to explicitly investigate whether these predictions manifest as neural activations of the expected bottom-up signal, i.e. tone, before the actual sensory input reaches the brain (15). Specifically, we sought to determine if, during sleep, the brain can pre-activate low-level features of an expected stimulus.

Participants (n=34) passively listened to tone sequences consisting of four simple 100 ms tones (low to high pitch; Figure 1a), while their brain activity was recorded via simultaneous EEG and MEG. Tone sequences were continuously presented for 20 minutes during wakefulness and during a 2.5-hour afternoon nap. We manipulated the transitional probabilities of the tones (Figures 1b, c), creating sequences in which predicting the next tone was either impossible (random sequences) or possible (predictable sequences). Tones were presented at a constant rate of 330 ms (3 Hz), thus establishing strong temporal predictions. Participants were neither instructed to pay attention to the tones nor informed about the transitional rules. Tone sequences were continuously presented for 20 minutes during wakefulness and during a 2.5-hour afternoon nap. All participants managed to fall asleep in the MEG scanner. Of these, 34 participants entered N1 sleep; 30 reached N2 sleep; 10 reached N3 sleep, and 6 reached REM sleep.

**Figure 1.**
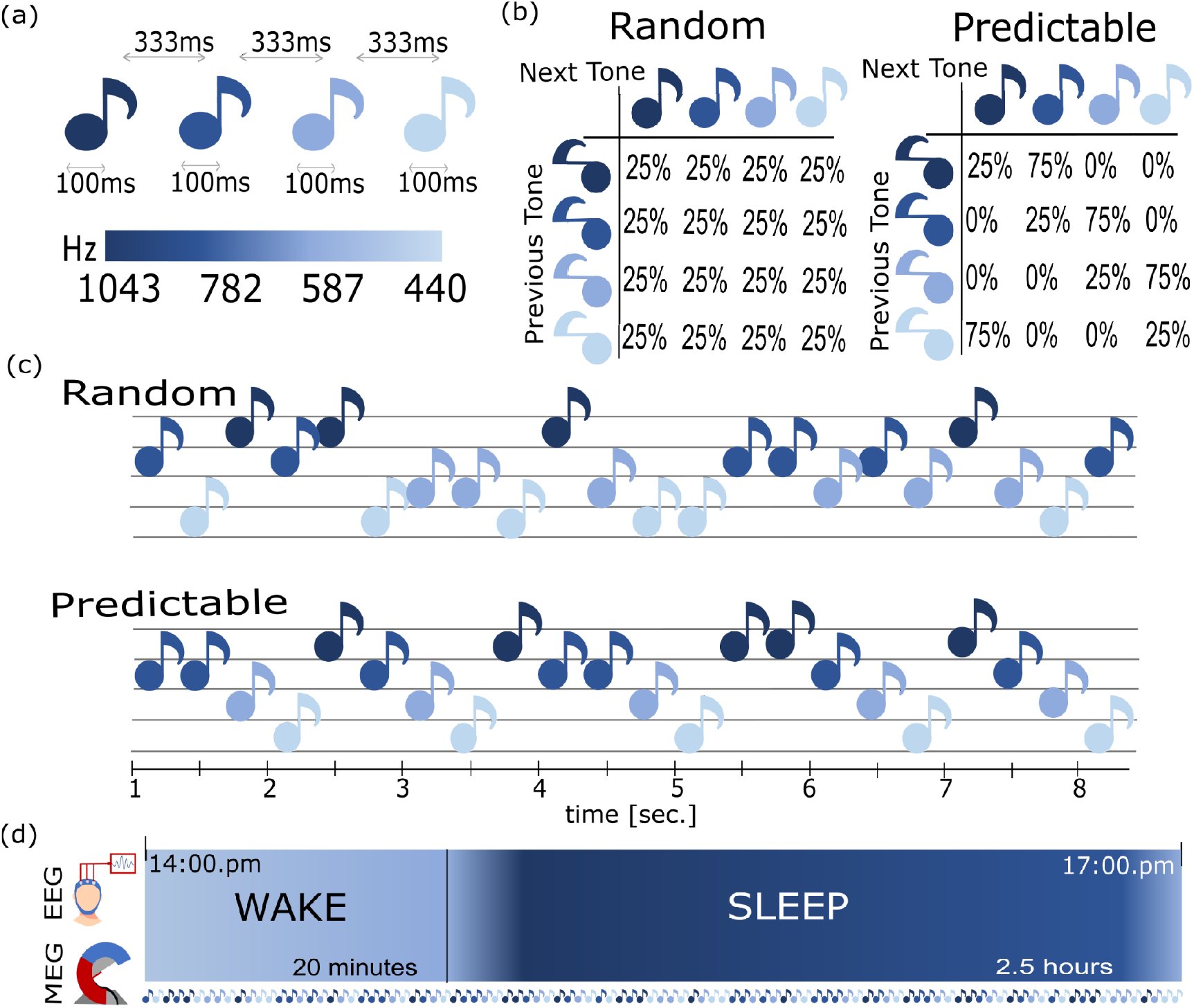
Experimental design and sleep protocol. **(a)** Four tones composed of four frequencies (low to high pitch: 440 Hz, B: 587 Hz, C: 782 Hz, and D: 1,043 Hz) were presented in a sequence at a constant presentation rate of 3 Hz (Inter-Stimulus-Interval 330 ms) for 100 ms each. **(b)** The transitional probabilities between the tones were manipulated creating random (25% chance of transition to either one of the four tones) and predictable (75% probability of transition to the next higher frequency tone) sequences that prevented or allowed the formation of feature-specific expectations (expectations about the frequency of the tone), respectively. As a result **(c)**, predictable tone sequences consisted of a reoccurring ascending tone progression (A to D) with few tone repetitions, whereas this was not the case for the random tone sequences. **(d)** While EEG and MEG were recorded simultaneously, participants passively listened to 5-minute blocks of random and ordered tone sequences during wakefulness, as well as during a 2.5-hour nap, untold of the underlying tone-sequence rules.

### The processing of subtle low-level stimulus features persists in N1 and N2 sleep

Having observed a robust response of auditory regions to sounds in wakefulness and sleep (see supplementary figure S1 for auditory event-field-potentials, ERFs), we first wanted to establish whether there are indeed unique neural activations that carry information about the carrier frequency of each tone. This is critical, because pure tones elicit highly correlated neural activations at the macroscopic level (16), localised in a small space within the auditory cortex (17). To that end, we used the single-trial MEG time-series data from the random tone sequences to train a linear discriminant analysis (LDA) classifier to distinguish the four different tones at each time point. As illustrated in Figure 2a, we observed significant above-chance (>25%) decoding accuracies, not only in wakefulness but also in N1 and N2 sleep. During wakefulness, a cluster-based permutation revealed two significant clusters (pcluster < .001, d=0.91) from 40 to 370 ms, and from 420 to 510 ms (pcluster_2_=.002, d=0.65). This sustained stimulus representation could reflect a memory trace that persists even after the onset of the next stimulus (at 330 ms in Figure 2). Such activation is crucial for forming associations across events, thus enabling the encoding of the underlying statistical regularities (4). Interestingly, a potential reactivation of the sensory input occurs at approximately 400 to 500 ms (cf. Figure 2), reflected in a transient increase in decoding accuracy.

**Figure 2.**
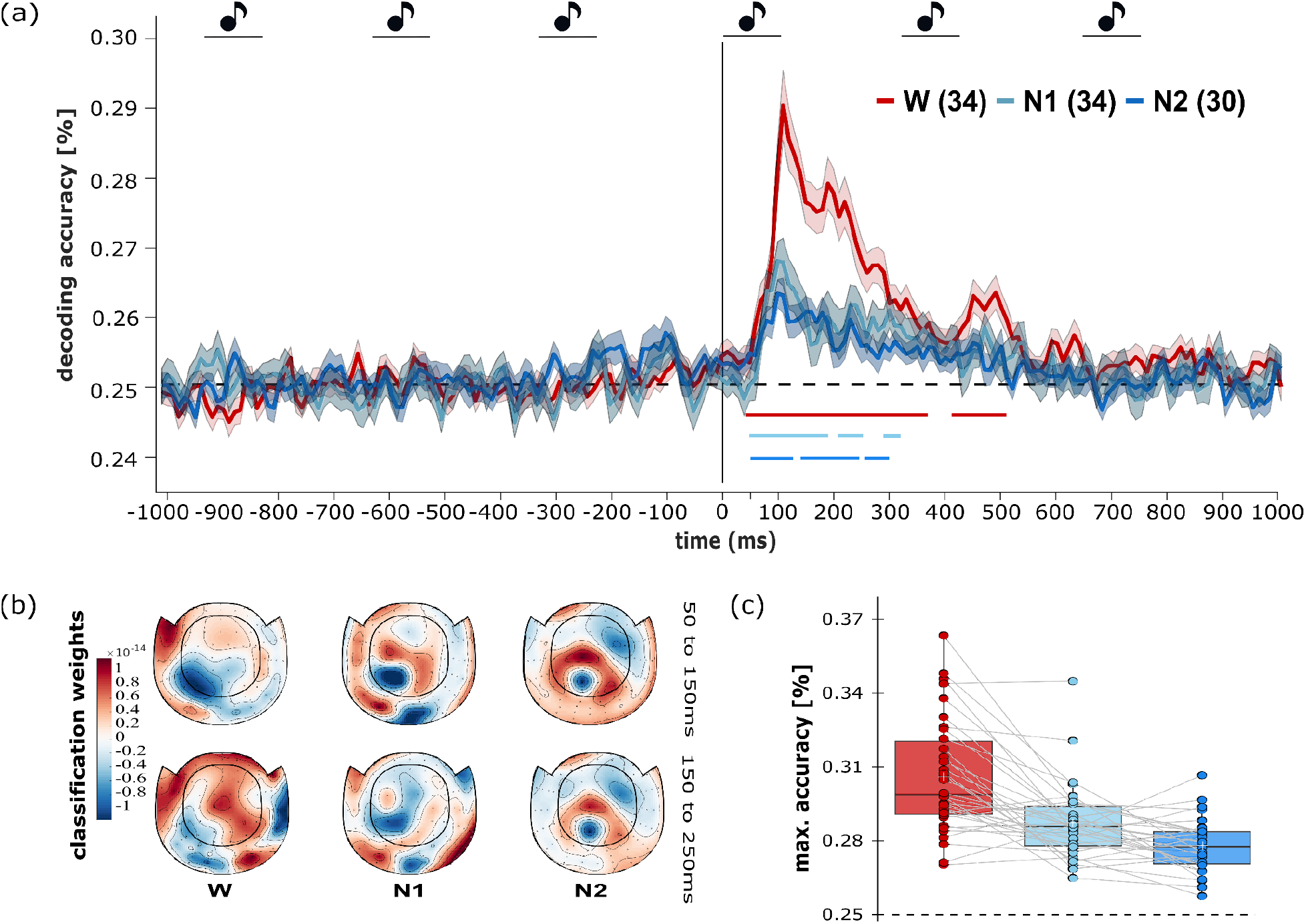
Decoding of low-level stimulus properties across time in wakefulness and sleep. **(a)** Significantly above-chance decoding accuracies shortly after stimulus presentation (0-100 ms), after training and testing only on magnetometer-derived single-trials of random tones. Coloured shades reflect the standard error of the classification accuracy. The dashed horizontal line marks the chance level at 25%, while the solid-coloured horizontal lines mark significant time clusters using cluster-based permutation analysis. Note that in wakefulness the tones could be decoded up to 550 ms after stimulus onset, replicating the findings of Demarchi et al. (2019). **(b)** Sensor-level normalised classification weights in wakefulness and sleep, separately for first and second classification accuracy peaks, as observed in wakefulness, N1 sleep and N2 sleep. **(c)** Boxplots depicting participants’ maximum classification decoding accuracy during the inter-stimulus interval (0 to 300 ms).

In N1 sleep, we observed three significant clusters: from 50 to 180 ms (pcluster_1_ < .001, d=0.57), from 200 to 250 ms (pcluster_2=_.01, d=0.47), and from 290 to 310 ms (pcluster_3_=.02, d=0.59). The tones were also distinguishable in N2 sleep, during which we observed four significant clusters: from 50 to 120 ms (pcluster_1_ <.001, d=0.61), 130 to 240 ms (pcluster_2_ <.001, d=0.71), 260 to 300 ms (pcluster_3_=.02, d=0.49), 340 to 390 ms (pcluster_4_=.03, d=0.50). These results suggest that even subtle changes in tone pitch are reflected in unique neural activations in both wakefulness and sleep.

For classification in wakefulness, left temporal channels contributed the most to the classification of tone features in the early-processing window (50-150 ms: Figure 2b-upper), while at later stages of the processing stream (150-250 ms: Figure 2b-lower), we observed a more widespread contribution of channels when the representations arguably become more abstract (Figure 2b-lower). For sleep, we did not observe such a pattern, finding instead that central channels were weighted as most informative (Figure 2b).

To compare the participant’s highest decoding accuracy in a given state, we computed the maximum classification accuracy in the post-stimulus-onset interval (0 to 330 ms) per participant and state. Classification accuracies were significantly different (F(2, 58)=28.7, p<.001) among wakefulness (M=30.3%, SD=2.1%), N1 (M=28.6%, SD=1.6%) and N2 (M=27.8%, SD=1.1%) sleep, with higher decoding accuracies in wakefulness than in N1 sleep (t(33)=3.63, p=.001) and N2 sleep (t(29)=3.63, p=.001), while decoding accuracies in N1 sleep were higher than in N2 sleep (t(29)=3.63, p=.001), cf. Figure 2c. These results suggest that compared to wakefulness, decoding accuracies in N1 and N2 sleep are significantly reduced, suggesting that feature-specific neural activations become less distinct.

### Sleep alters the temporal dynamics of auditory neural activations

We found evidence for the ability of the brain to carry feature-specific information in wakefulness and sleep. Next, we aimed to explore how such feature-specific neural activations dynamically change over time: how temporarily available they are (18). To that end, we used a time-generalization approach, in which we trained and tested a classifier at different time points. This approach enabled us to determine if neural activations at time t are also detectable at another time t’.

Time-generalization decoding revealed significant clusters after stimulus onset in all states: wakefulness (pcluster <.001, d=0.4), N1 (pcluster1 <.001, d=0.12) and N2 (pcluster1 <.001, d=0.22). We observed a decreasing size of the significant cluster along the diagonal from wakefulness to sleep, suggesting a restriction on how temporally accessible the feature-specific neural activations are as individuals progress from wakefulness to sleep (cf. Figure 3). This suggests that in sleep, frequency-specific neural activations are sequentially activated and are less temporarily available.

**Figure 3.**
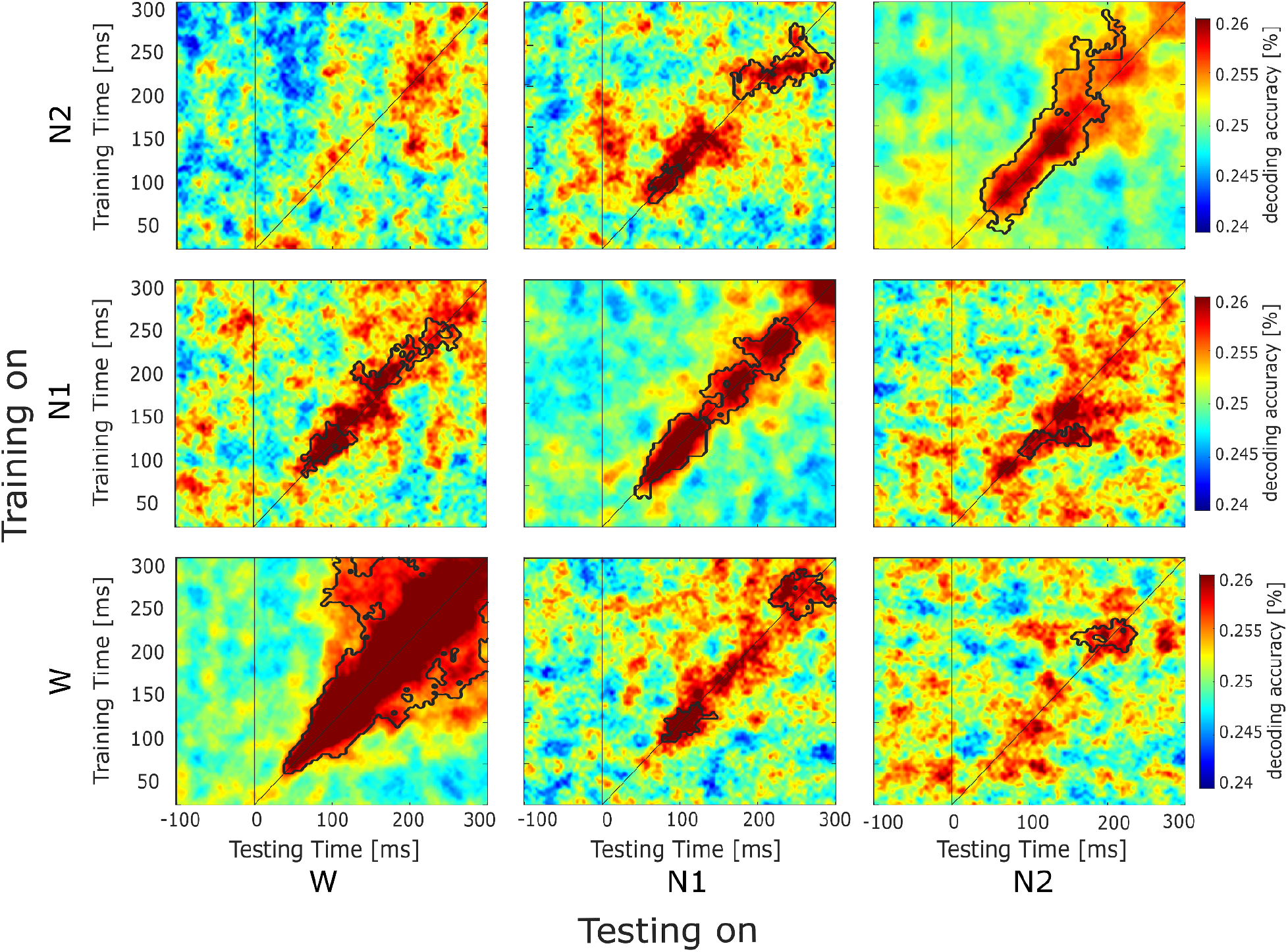
Within- and cross-state temporal-generalization matrices. Decoding accuracies of classifiers trained (y-axis) and tested (x-axis) separately on each time point after stimulus onset (0 to 300 ms). Interestingly, feature-specific neural activations in sleep are short-lived and less widespread over time, suggesting that they are less available over time. Cross-state decoding accuracies suggest there is some similarity between the feature-specific neural representations across the wakefulness, N1 and N2 stages. Black frames illustrate the time points at which the cluster-based permutation test yields significant results when compared to the chance distribution of shuffled labels.

To examine the similarity of the feature-specific neural representations between wakefulness and sleep, we performed cross-state decoding. Specifically, we trained an LDA classifier to distinguish among the tones in one state (e.g. wakefulness) and then tested its performance in another state (e.g. N2). This approach allowed us to assess whether the neural activations associated with tone features were consistent across states (Figure 3).

We observed significant clusters when the classifier was trained on wakefulness and tested on N1 (pcluster_1_ <.001, d=0.34, training time: 100-150 ms, testing time: 50-15 ms; pcluster_2_=.02, d=0.35, training time: 230-400 ms, testing time: 200-300 ms) and N2 sleep (pcluster=.01, d=0.35), cf. Figure 3. We also observed above-chance decoding accuracies when training the classifier on the N1 trial and testing it in wakefulness (pcluster_1_ <.001, d=0.28, training time: 100-150 ms, testing time: 50-150 ms; pcluster_2_=.01, d=0.26, training time: 200-250 ms, testing time: 100-250 ms) and N2 trials (pcluster <.001, d=0.3). Similarly, training on N2 revealed significant above-chance clusters when tested on N1 (pcluster_1_ <.001, d=0.30), but not in wakefulness trials. These results suggest that we can successfully decode the features of a tone in one state by using a classifier that has been trained in another state.

Altogether, these results demonstrate that while feature-specific neural activity is sustained during wakefulness, especially at later processing stages, during sleep, informative patterns do not generalize across time. However, neural activations during wakefulness can still be used to discriminate among the tones presented during N1 and N2 sleep, indicating that feature-specific neural activations share some similarities across wakefulness and sleep. This is most pronounced for later processing stages from 200 to 300 ms.

### Anticipatory pre-activations decline from wakefulness to sleep

An anticipatory effect of predictions should be reflected in an increase of feature-specific activations before the onset of an expected tone, i.e. pre-activation. To search for such pre-activations, we trained our classifier exclusively on post-stimulus (0 to 300 ms) data from the random tone sequences. This ensures that the classifier learns exclusively tone-specific activation patterns (i.e. post-stimulus window) and avoids any carryover effects from the high chance (75%) of encountering the same tone (i.e. random sequences) in the predictable sequence (see star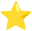Method). We then tested the performance of this classifier in the random and predictable pre-stimulus interval, hypothesizing that predictable tone sequences alone would lead to higher pre-stimulus decoding accuracies (4).

Indeed, in wakefulness, we observed significantly higher classification accuracies in predictable sequences than in random ones when the classifier was trained on the approximately 100 to 250 ms post-stimulus window and tested on the approximately -300 to -150 ms pre-stimulus window (pcluster< .001, d=0.52), replicating previous findings (4). This pre-activation suggests that stimulus-specific neural representations are activated before the actual presentation of the expected stimulus, providing evidence for anticipatory brain activity.

In addition, we found a significant post-stimulus cluster (*p*cluster=.01, d= 2.87), that involves activations from approximately 100 to 250 ms (cf-Figure 4). A pre-activation pattern similar to wakefulness was observed in N1 sleep (Figure 4-middle column) at approximately -300 to -200 ms (*p*cluster=.023, d=0.31). No significant pre-stimulus clusters were identified in N2 sleep when compared to the other two conditions (Figure 4 left column). Interestingly, the absence of pre-activations in N2 was observed while post-stimulus on-diagonal decoding was above chance, demonstrating preserved feature-specific auditory processing.

**Figure 4.**
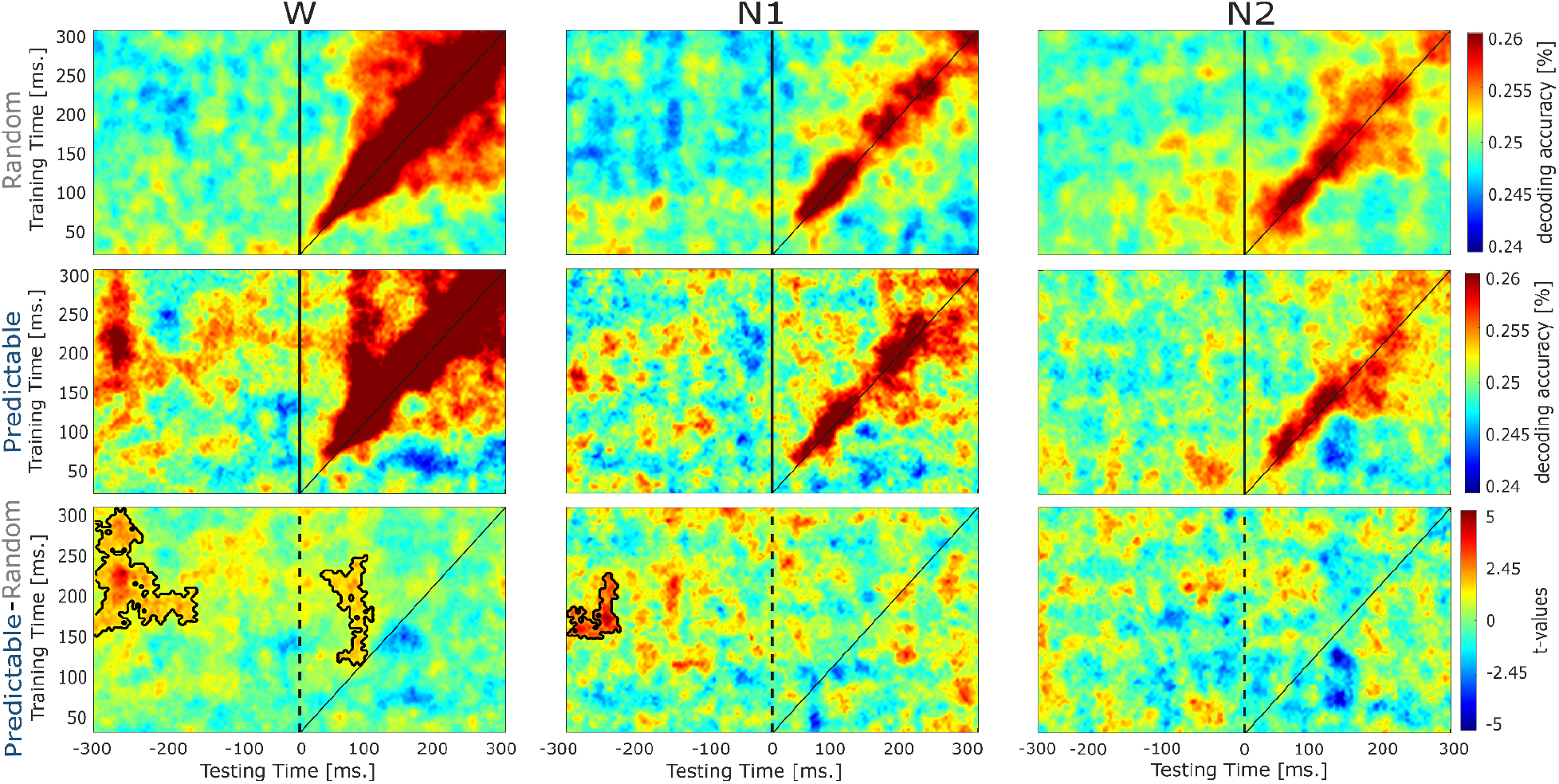
Pre-activations of low-level features of the expected stimulus in wakefulness and sleep. Raw grand average time-generalization decoding accuracies training on single-trial raw random tone activations and tested on (top) random and (middle) ordered tones. Paired samples t-values contrasting ordered to random tones (bottom) using a non-parametric cluster-based permutation. Black frames illustrate the time points where the non-parametric cluster-based permutation test yields significant results. Note that in wakefulness, the brain pre-activates the low-level stimulus properties when they can be predicted (i.e., in the ordered condition), as is the case for N1 sleep. However, during N2 sleep, there is no evidence of such pre-activations, although the feature-specific on-diagonal decoding leads to above-chance decoding accuracies.

## DISCUSSION

Our study reveals that the brain does not actively engage in predicting the features of upcoming stimuli during sleep (Figure 4) and that neural representations in sleep appear to be less distinct (Figure 2) and sustained (Figure 3), possibly hindering the tracking and extraction of embedded associations. These results expand our understanding of feature-specific predictions in sleep and demonstrate the effectiveness of MVPA for studying human cognition covertly, as required in unresponsive states, such as sleep.

Since sleep is a reversible state, the brain is not completely disconnected from the environment, continuously processing as it does upcoming sensory input (19). Our results demonstrate that the auditory cortex, for example, reacts to upcoming stimuli (supplementary Figure 1: ERFs) and even encodes details of sensory inputs, as shown in Figure 2. Here we reveal the extent of such processing by showing that even subtle changes in auditory stimuli (i.e., changes in carrier frequency) with no embedded higher cognitive properties, such as semantics, self-relevance (20), or emotion (21,22), and without explicit allocation of attention upon encoding, can be differentiated in sleep.

Observing sustained above-chance decoding accuracies in sleep that surpass the stimulus presentation duration (beyond 100 ms) suggests that the decodable information is beyond simple sensory encoding because the low-level stimulus properties are going to a higher representational space. However, the transformation of sensory inputs to percepts is compromised during sleep, given that even advanced methods such as MVPA — which is far less susceptible to the endogenous “noise” that sleep introduces than are contemporary methods (e.g., event-related responses) — struggle to identify the unique feature-specific neural patterns in sleep, a contrast to the effectiveness of these methods applied to wakefulness. The drop in classification accuracies in sleep can be explained by the reduction in the feature-specific information available to aid the classifier in distinguishing the four tones for successful decoding. This is in line with invasive local field potential evidence from animal studies, which suggests that the information-transfer ratio between the output and input of visual thalamocortical relay neurons gradually decreases from wakefulness to deeper sleep stages (12). Such reduction might explain the less temporarily available neural activations (cf. Figure 3), suggesting that feature-specific neural activations are short-lived and potentially restricted in their cortical propagation during sleep (Razi et al., 2023), albeit observed similar activations of auditory hubs, upon auditory stimulation (Figure S1). Together, these findings point towards an effective relay of sensory input to primary cortices during sleep, with some loss of low-level stimulus features (23), as well as loss of information integration at later cortical-process stages (10,11).

Many theoretical accounts propose that perception is understood as a process of inference, relying on internal models that generate automatic predictions (generative models) about the most plausible sensory interpretation, based on prior information (3). These predictions manifest as neural activations of the expected bottom-up signal before the actual sensory input reaches the brain (15) and have been recently empirically demonstrated in wakefulness in the visual (1) and auditory (4,24) modalities, even in the absence of the expected bottom-up input (7), i.e. when the actual predicted stimulus is omitted. Predictions in sleep have been extensively studied using either simple event-related responses (14,19), or more recently, continuous auditory stimulation and frequency tagging (7–9). For the first time, to our knowledge, we explicitly searched for evidence of feature-specific pre-activations and concluded that anticipatory processing ceases as the brain transits from wakefulness to sleep. The brain’s inability to represent statistical associations during sleep, in the presence of intact preserved sensory processing reported here, is in line with recent studies investigating brain entrainment to statistical regularities (8,9), as well as with studies using invasive methods that report disruption of feedback processing, a prerequisite for predictive processing, challenging findings that show differential responses to expected versus unexpected stimuli (5,6,11). These approaches are limited, as it is difficult to conclude whether the disruption of predictive processes observed in sleep arises from fundamental changes in underlying perceptual or predictive computations from those operating in wakefulness, or whether they arise rather from sleep-related modifications of brain electrophysiology (25). Here, we used a highly sensitive method, MVPA, which should be less affected by the uninformative modifications in sleep-related electrophysiology, since it capitalizes on unique brain patterns that drive higher classification. Surpassing the limits of previous attempts and utilizing non-invasive methods such as single-trial MEG responses, we conclude that the brain ceases to engage in predictive processing during sleep. This may be due to a failure to sustain a memory trace that would enable the extraction of sensory associations over time.

The extent to which prediction would prevail in sleep after extensive exposure to predictable tone sequences, as well as the explicit encoding of the underlying tone-sequence structure (i.e., making participants aware of the underlying rules), are fruitful avenues for future research. Unfortunately, the number of participants to reach N3 and REM sleep did not allow for robust statistical analysis; we provide, however, a descriptive overview of the results in the supplementary material (see Figure S2).

In sum, we extend previous studies by using a sensitive methodology that allows for searching for feature-specific pre-activations in sleep. Our results suggest that the brain ceases to engage in the active prediction of sensory inputs in sleep, even as basic perceptual processes continue to operate.

## Declaration of generative AI and AI-assisted technologies in the writing process

During the preparation of this work, the author(s) used ChatGPT v4.0 to improve readability. After using this tool/service, the author(s) reviewed and edited the content as needed and take(s) full responsibility for the content of the publication

## Star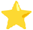Methods

### Participants & Sleep Descriptives

We recruited 37 healthy participants for the present study. The data of three participants were discarded due to technical issues with the recordings. Thus, the final dataset consisted of 34 participants, the majority of which were male (females=9), with a mean age of 27.4 years (SD=5.8, range=20 - 41). We explicitly looked for healthy participants with no previous psychiatric or neurological disorders, including any sleep problems. As lying down without moving in the MEG can be uncomfortable, we included only those participants who reported that they can fall asleep quickly and can maintain sleep while sleeping on their backs. All participants entered N1 sleep (N=34), while 30 participants reached N2 sleep, and 14, N3 sleep. Six participants even reached REM sleep in the MEG scanner. Because too few participants reached N3 and REM sleep, the results in these states are presented in the supplementary materials (Figure S2). The experimental protocol was approved by the ethics committee of the University of Salzburg, and all participants provided written consent.

### Experimental Design and Stimuli

We adjusted the experimental design from Demarchi and colleagues (4). Having mounted participants with an EEG cap and five head-position-indicator (HPI) coils, then having marked anatomical landmarks and digitized their head model, we placed them in the shielded MEG room in the supine position. We made sure that participants were lying comfortably in the scanner and used additional pads to ensure utmost comfort throughout the experiment, while minimizing head movements. After a five-minute resting-state recording, we determined each participant’s hearing threshold using an auditory staircase. We manually adjusted the volume until participants indicated a comfortable volume level that we could use during sleep. The sound loudness was kept constant during both wakefulness and sleep, as the sound intensity can influence the evoked brain responses (26). We presented the participants with tone sequences (i.e., tones presented one after the other), composed of four different pure (sinusoidal) low- and high-pitch tones (A: 440 Hz, B: 587 Hz, C: 782 Hz, and D: 1,043 Hz), at a constant 3 Hz presentation rate (i.e., every 333 ms). Each tone lasted 100 ms (5 ms linear fade in/out; see Figure 1). In wakefulness, participants listened to four blocks of tone sequences, each containing 1000 stimuli, with each block lasting about 5 minutes. While within each block the overall number of presentations for each tone was equal, we parametrically manipulated their transitional probabilities, creating sequences of random and ordered tones.

In detail, the random sequences were characterized by equal transitional probabilities from one sound to another (25% chance for one of the four tones to be presented), making it impossible to predict which of the four tones would next appear (see Figure 1b, c). In contrast, in ordered tone sequences, the probability that a tone would be followed by the next-higher-in-pitch tone was quite high (75%), thus allowing the prediction that a higher-pitch tone would follow. As in the random condition, there was a 25% chance of tone repetition (see Figure 1b) to ensure equal carryover effects across conditions. To decrease the possibility of participants sleeping during wakefulness stimulation, participants wore in-house-built prism glasses that enabled them to watch a movie on a projector screen without sounds or language-embedded scenes (i.e., underwater wildlife), while passively listening to the tone sequences. To further decrease the probability that the participants would fall asleep during wakefulness stimulation, we checked the state of the participants between blocks and reminded them of the number of remaining blocks. After listening to four blocks in wakefulness, the lights were dimmed, and participants were asked to remove their prism glasses and to take a nap, during which the tone sequences would continue to play in the background. To increase the probability that the participants would fall asleep in the scanner, and thus sleep propensity, we instructed participants to sleep two hours less than usual the night before testing. In wakefulness, the tone-sequence blocks switched from random to ordered (R-O-R-O or O-R-O-R), while in sleep, the order of the tone sequences was pseudo-randomized, with no more than two of the same tone sequence occurring in succession. Participants were awakened after 2.5 hours in the scanner, and the experiment was terminated (Figure 1d). At the end of the experiment, 24 out of 34 participants were asked if they had noticed that there were two tone sequences, but only three answered affirmatively. The experiment was programmed in Matlab 9.1 (The MathWorks, Natick, Massachusetts, USA), using the o-PTB toolbox (27).

### Signal Acquisitions

We used the Elekta Neuromag Triux (Elekta Oy, Finland) system to record whole-head brain magnetic signals at a sampling rate of 1000 Hz, with 0.1–330 Hz hardware filters, in a standard, passive magnetically shielded room (AK3b, Vacuumschmelze, Germany). Signals were captured by 102 magnetometers and 204 orthogonally placed planar gradiometers at 102 different positions. We used a Polhemus FASTTRAK digitizer to sample the anatomical landmarks (nasion and left/right pre-auricular points), the HPI locations, and around 300 head-shape points. In addition, for accurately measuring sleep, we recorded polysomnography data using 16 single-channel EEG electrodes (F3, Fz, F4, T3, T4, T5, T6, C3, Cz, C4, Pz, O1, Oz, O2, and Mastoid electrodes A1 and A2), left and right horizontal EOG channels, two chin EMG channels and an ECG channel. Upon recording, we visually examined the channel impedance below 5ω and adjusted until this criterion was met.

### MEG Data Pre-processing

We applied a signal-space-separation algorithm implemented in the Maxfilter program (version 2.2.15) to remove external noise (mainly 16.6 Hz, and 50 Hz, plus harmonics) and to realign data to a common standard head position (trans default Maxfilter parameter). We applied a high-pass filter (0.1 Hz) and defined 2-second trials (with 1-second pre- and post-stimulus intervals) for both wakefulness and sleep. We conducted a semi-automatic independent component analysis (ICA) to visually identify and remove eye blinks and heartbeat during wakefulness and sleep. In a semi-automatic manner, we identified and excluded trials containing (jump) artifacts. Following Demarchi et al. (2019), we included only the magnetometers, while the EEG data was used exclusively for sleep staging and was not analyzed further. Finally, the epoched data were high-pass filtered at 0.1 Hz (Butterworth IIR filter) and downsampled to 250 Hz. The data was preprocessed using the Fieldtrip toolbox in Matlab (28).

### Sleep staging

To extract sleep stages, the polysomnography data were pre-processed following the guidelines of the American Association for Sleep Medicine (AASM;25).

Specifically, the EEG and EOG data were preprocessed using a 40-Hz low-pass filter and a 0.1 high-pass filter, while the EMG data was band-pass filtered between 10 and 100 Hz. We then performed automatic sleep-stage classification using the Alice Sleepware G3 Software (Philips Respironics, Murrysville, PA, USA) which characterizes sleep stages according to the AASM criteria and has been validated against sleep staging by human experts (30), displaying human-level sleep-stage classification capabilities.

### Multivariate Pattern Analysis (MVPA)

MVPA was performed in Matlab (Matlab 2012 - version R2020a) using the MVPA-Light toolbox (31). The preprocessed data were further filtered using a low-pass filter at 30Hz. A Multiclass Linear Discriminant Analysis (LDA) classifier was used on single-trial sensor-level data (102 magnetometers; features) to discriminate between the four tones (classes), with the chance level at 25%. Throughout this manuscript, we have used only the random trials for training, as they are described by equal transitional probabilities and, thus, equal low-level stimulus predictions. We controlled any further imbalances by equalizing the number of each of the four tones in all analyses. The classification was performed at each time point, using 5-fold cross-validation. To examine whether there are unique and distinguishable neural activations corresponding to the four tones reflected in above-chance decoding, we used across-time decoding (training and testing at the same time point, e.g., training a classifier at 100 ms and testing at 100 ms). To examine how those activations dynamically evolve over time, instead of applying a different classifier at each time point, the classifier trained at time t (e.g., 100) can be tested on its ability to generalize to time t’ (e.g., 150). This approach — training and testing at different time points — is referred to as time-generalization decoding and is described in detail by King & Dehaene (18). In the following subsection, we provide further details for each of our analyses:

#### Examining the processing of low-stimulus features

We first performed across-time decoding in wakefulness and sleep to determine the extent to which the low-level stimulus features are distinguishable. Using cross-validation, we trained and tested a classifier on the maximum number of random trials found in each state to maximize the available data for learning. On average, per participant, there were 1,879 (SD=235; min 975) random trials in wakefulness, 2,472 (SD=1,970) in N1 (min=284), and 3,820 trials in N2 (SD=2,940; min=497), with N2 having statistically the highest number of trials (p<.001). To compare the participant’s highest decoding accuracy in a given state, we extracted the maximum across-time classification accuracy in the post-stimulus-onset interval (0 to 330 ms) per participant and state. Furthermore, we used time-generalization decoding to explore how the neural activations change over time within each state separately (i.e., within-state decoding: training in N1 and testing in N1), but also to see how similar the neural activations were between states by training and testing in different states (i.e., between-state decoding: training in N1 and testing in N2). To ensure that differences in decoding accuracies did not result from different amounts of training data per state, we made sure the classifier was trained and tested on an equal number of trials across states. Thus, we identified the minimum number of trials across states for each participant and randomly selected the same number of trials for each state. In the case that fewer than 200 trials were presented in a state (that is, less than 1 minute of stimulation and 50 trials per tone), the state was not included in the analysis. In this analysis, there were, on average, 1,180 (SD=546, range=284-1,990) trials per participant that were used for training the classifier. To visualize which channels contributed to classification, we extracted the balanced classifier weights (i.e., the product of the classifier weights and the data covariance matrix as suggested in Haufe et al. (32). For visualization purposes, we z-transformed the classifier weights for each state without applying baseline normalization. Finally, we explored which of the tones drove the classification accuracies by decoding the low-versus high-pitch tones and the middle-pitch tones separately for each state.

#### Searching for feature-specific stimulus pre-activations

To investigate whether the neural response to a stimulus (e.g, B) is present before its actual presentation (pre-activation) as a result of being predicted by a preceding stimulus (4), we examined stimulus pre-activations by looking at the pre-stimulus decodability of the upcoming stimulus using time-generalization decoding, i.e. the ability to decode stimulus-specific neural activity in the pre-stimulus window using neural activity immediately following stimulus presentation. We used only the random tones to train a multiclass linear classifier to avoid any potential carryover effects (i.e. the classifier picking up information from preceding trials if transitions are predictable). There were no extra preprocessing steps.

### Statistical Analysis

Throughout our analyses, we used a nonparametric cluster-based permutation to determine whether the decoding accuracies for the random and predictable classification accuracies can be drawn by a similar distribution while controlling for the multiple comparisons (33). The data were partitioned 10,000 times, and the decoding accuracies were compared against a chance distribution by shuffling the labels at each time point using a one-tailed (alpha=.025) between-sample t-test. When statistically comparing directly how the similarity between tones (i.e, low-vs. high-pitch and middle tones) drives decoding accuracies, we performed a cluster-based permutation using a dependent-samples t-test (two-tailed; alpha=.05). A similar approach was used when statistically comparing the predictable to the random decoding accuracies. There was no cluster permutation analysis in which there was a selection of time windows (i.e., the displayed time windows are the ones that were used in the cluster-based permutation without any *a priori* time-window selection). Following guidelines about good practice for conducting and reporting MEG research, we report the average Cohen’s d for each significant cluster, based on which we rejected H0 (34). We statistically analysed the maximum per participant and state classification accuracies using Analysis of Variance (ANOVA) and followed up with a dependent-samples t-test.

#### Code accessibility

The code used for data analysis is available at the first author’s Gitlab repository: https://gitlab.com/PavlosTopalidis/sleepmarkov.

## Author contributions

P.T. and L.R. collected the data; P.T., N.W., and M.S. designed the experiment; P.T. wrote the manuscript; P.T. and J.S. implemented the experimental protocol; P.T. conducted data analysis; N.W., M.A., J.S., and L.R. edited the manuscript; N.W., M.S., and M.A. supervised the work.

## Acknowledgements

Pavlos Topalidis was supported by the Doctoral College “Imaging the Mind” (FWF, Austrian Science Fund: W 1233-B).

## Supplementary

**Figure S1.**
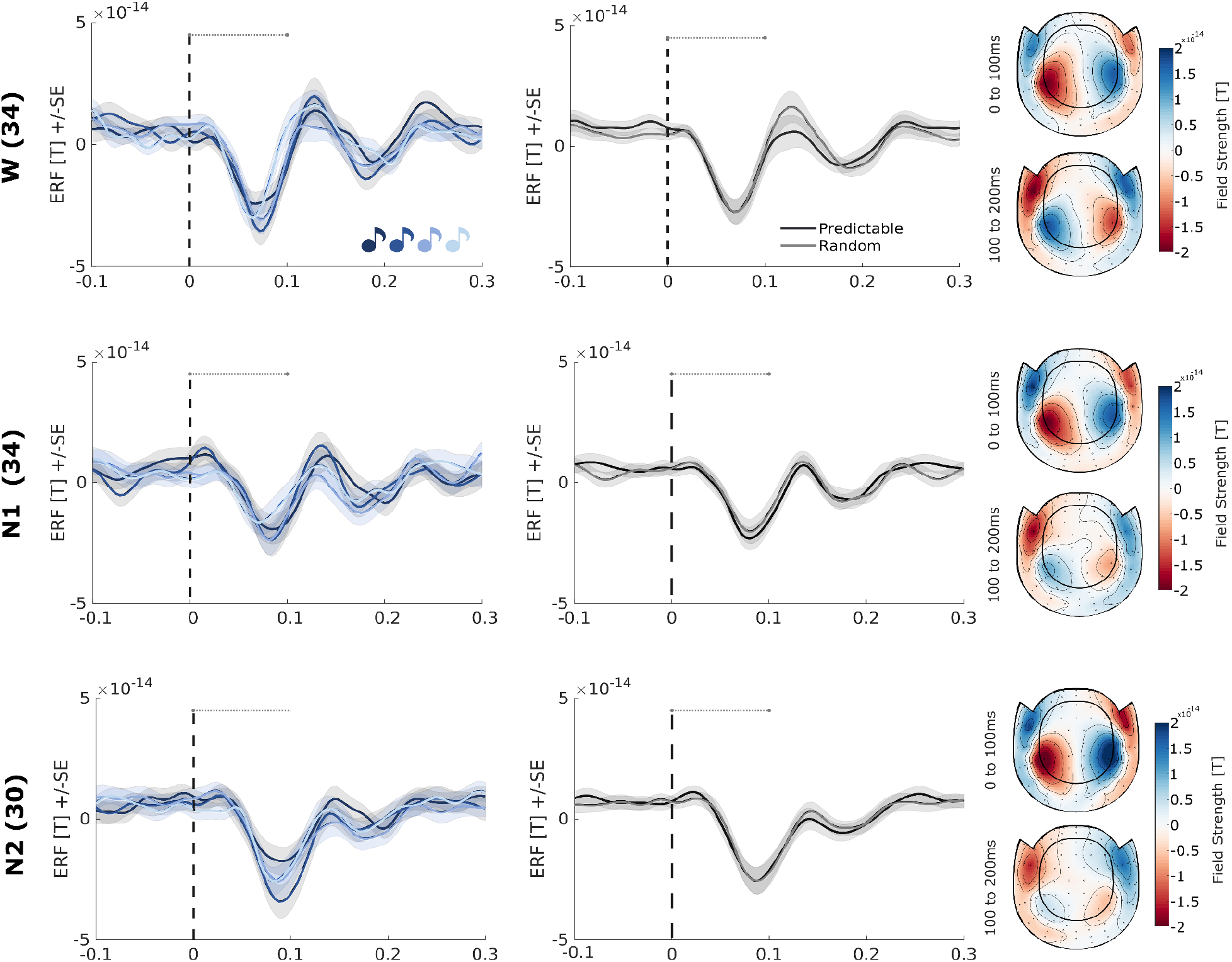
Grand averaged Event Related Fields (ERFs) of magnetometer auditory channels per tone and regularity locked to the stimulus onset (0 ms), in wakefulness, N1 and N2 sleep. Only the four random tones were used to display the ERFs (left) with their topographic activations (right) from 0 to 100 ms and 100 to 200 ms (right). The grand average ERFs of the predictable (black) and random tones (gray) tones are also displayed. The horizontal dotted line marks the stimulus duration. Note that the auditory channels react upon stimulation in N1 and N2 sleep, while the amplitude of such responses slightly decreases.

**Figure S2.**
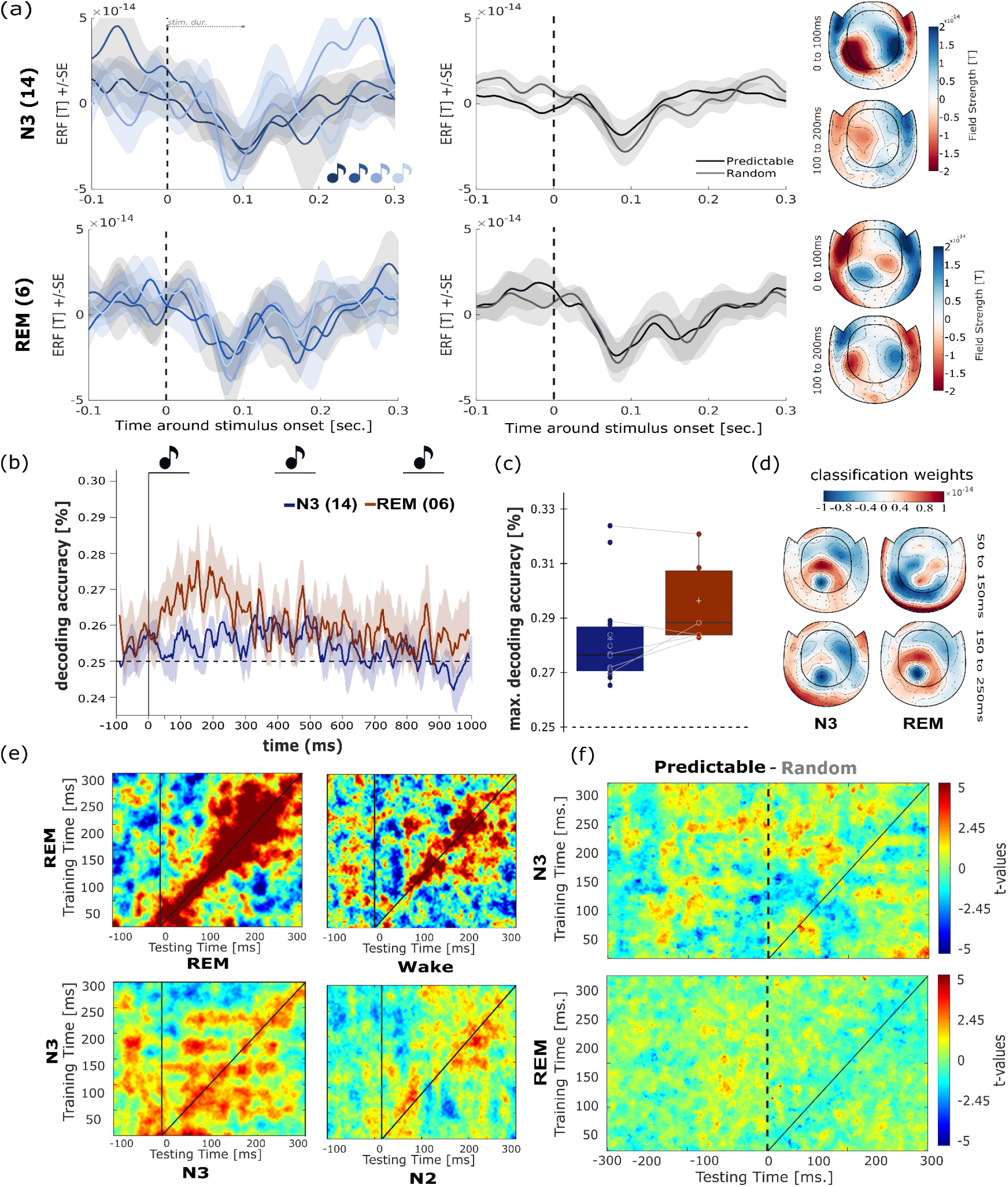
Descriptive results in N3 and Rapid Eye Movement (REM) Sleep. **(a)** Grand averaged Event Related Fields (ERFs) of auditory magnetometers per tone and regularity locked to the stimulus onset (0 ms; vertical dashed line). It appears that the expected auditory responses at 100 ms are observed in N3 and REM sleep. **(b)** Decoding of low-level stimulus properties across time appears to be descriptively above chance in REM sleep, albeit in a few participants (N=6). **(c**,**d)** When looking at the maximum decoding accuracies for the participants who achieved both N3 and REM sleep, we descriptively observe that higher decoding accuracies are obtained in REM sleep, with posterior and central channels contributing to this classification. **(e)** High on-diagonal decoding accuracies are observed in REM sleep, even when testing on wakefulness trials, implying similarity in their neural representations. Time generalization in N3 sleep displays an oscillatory pattern possibly reflecting the underlying N3 sleep physiology while using the N2 trials as a testing set suggests some high on-diagonal accuracies. **(f)** T descriptive level there was no higher decoding accuracies at the prestimulus interval in N3 and REM sleep (here displayed as t values of predictable minus random accuracies).

